# Impact of Cannabinoid Type 1 Receptor Modulation on Risky Decision-Making

**DOI:** 10.1101/2020.03.13.990721

**Authors:** Timothy G. Freels, Anna E. Liley, Daniel B. K. Gabriel, Nicholas W. Simon

## Abstract

Recent changes in policy regarding cannabis in the U.S. have been accompanied by an increase in the prevalence of cannabis use and a reduction in the perceived harms associated with consumption. However, little is understood regarding the effects of cannabinoids on cognitive processes. Given that deficient risk-taking is commonly observed in individuals suffering from substance use disorders (SUDs), we assessed the impact of manipulating cannabinoid type 1 receptors (CB_1_Rs; the primary target for Δ^9^-tetrahydrocannabinol in the brain) on punishment-based risk-taking using the risky decision-making task (RDT) in male Long-Evans rats. The RDT measures preference for small, safe rewards over large, risky rewards associated with an escalating chance of foot shock. Systemic bidirectional CB_1_R manipulation with a CB_1_R agonist, CB_1_R antagonist, and FAAH inhibitor (which increases overall endocannabinoid tone) did not alter overt risk-taking in the RDT. Interestingly, direct CB_1_R agonism, but not indirect CB_1_R stimulation or CB_1_R blockade, resulted in reduction in latency to make risky choices while not altering safe choice latency. Our findings suggest that CB_1_R activation expedites engagement in punishment based risk-taking without affecting overall preference for risky vs. safe options. This indicates that risk preference and rate of deliberation for risk-taking are influenced by distinct neural substrates, an important consideration for development of precise treatments targeting the aberrant risk-taking typical of SUD symptomology.

## INTRODUCTION

Cannabis has been recognized as one of the most commonly used psychoactive substances in the U.S. (National Institute on Drug Abuse, 2019). Widespread shifts toward more permissive cannabis-related legislation have been accompanied by a marked decrease in the perceived harms of cannabis use and concomitant increase in self-reported cannabis consumption among U.S. citizens (Compton et al., 2016). Additionally, sales of cannabis products containing high concentrations of the primary psychoactive phytocannabinoid Δ^9^-tetrahydrocannabinol (THC) have also become more prominent in legal markets (e.g., Washington state; Smart et al., 2017). As such, there is legitimate concern regarding the increase in availability of recreational cannabis given that the effects of cannabinoids on cognition are not fully understood.

A particularly critical component of cognition is risky decision-making, the process of determining whether to seek rewards accompanied by adverse consequences over safer (but often less gratifying) options (Simon et al., 2011; Gabriel et al., 2019). In humans, increased risk-taking stemming from overvaluation of reward and diminished sensitivity to adverse outcomes is common among people suffering from substance use disorders (SUDs), and promotes the onset and maintenance of drug use (Lane and Cherek, 2000; Kreek et al., 2005). Acute and chronic cannabis use increases risk-taking in human participants in laboratory models of risk-taking/gambling (Lane et al., 2005; Fridberg et al., 2010). Given that the cannabinoid type 1 receptor (CB_1_R) is the primary target for THC binding in the brain, it is possible that THC-induced CB_1_R activation might lead to increased risk-taking, which could promote problematic drug use.

Animal models of risk-taking such as the rodent gambling task have been used to assess the impact of CB_1_R modulation on risk preference to circumvent common extraneous variables encountered in human research. These studies have shown that administration of THC, synthetic CB_1_R agonists, or inhibitors of endocannabinoid catabolic enzymes has little impact on risk preference in the rat Gambling Task; although, some synthetic CB_1_R agonists have been shown to improve decision making in rats that initially exhibit suboptimal choice patterns (Gueye et al., 2016; Ferland et al., 2018). However, these studies utilized a modality of risky decision-making in which the reward and punishment each manipulated a single outcome; a risky choice may yield a large food pellet reward or remove access to that pellet reward. As such, it is still unclear how CB_1_R modulation might affect risky decision-making in which the reward and punishment are unrelated constructs. This is a critical distinction because the consequences of risk-taking are often distinct from the rewards; for example, substance use may be associated with legal ramifications such as incarceration or fines.

The Risky Decision-Making Task (RDT) addresses this by giving rats the choice between a small, safe food reward and a large, risky food reward with a varying chance of a mild foot shock (Simon and Setlow, 2012). Critically, performance in the RDT has been shown to reflect punishment-based risk-taking as an independent construct that is not influenced by differences in motivation, pain sensitivity, or anxiety-like behavior (Simon et al., 2011). Risk-taking in the RDT has also been shown to be predictive of behaviors and pharmacological factors associated with SUD such as impulsive action, psychostimulant self-administration, and nicotine sensitivity (Mitchell et al., 2014; Gabriel et al., 2019). Therefore, the RDT provides a useful means to assess the impact of CB_1_R manipulation on risk-taking and to investigate whether CB_1_R activation-induced modulation of risk preference might lead to problematic drug use.

Here, we utilized RDT with acute systemic administration of a battery of cannabinoid drugs to determine whether direct CB_1_R stimulation, enhancement of endocannabinoid tone, or CB_1_R blockade alter risky decision-making. Arachidonoyl-2-chloroethylamide (ACEA) was chosen to selectively stimulate CB_1_R receptors due to its high affinity for CB_1_Rs (Hillard et al., 1999). Arachidonoyl serotonin (AA-5-HT) was used to enhance endocannabinoid tone by both inhibiting the endocannabinoid catabolic enzyme fatty acid amide hydrolase (FAAH) and reducing off-site binding of endocannabinoids to transient receptor potential vanilloid type 1 channels (TRPV_1_s; Maione et al., 2007; de Novellis et al., 2008). CB_1_Rs were blocked using the inverse agonist rimonabant. Together, our experiments aimed to investigate how cannabinergic signaling might contribute to SUD symptomology by modulating risky decision-making.

## METHODS

### Subjects

Adult male Long-Evans rats (*N* = 19; Envigo) were pair-housed and maintained on a reverse 12:12 light/dark cycle with lights off at 08:00. To increase motivation for reward-seeking, rats were food restricted to 90% of free feeding body weight following a one week habituation period. If aggression or food dominance occurred, rat pairs were separated. Rats began training in the RDT at approximately 80 days of age. One cohort (*n* = 9) was used for ACEA and AA-5-HT experiments, and a second cohort (*n* = 10) was used for rimonabant experiments. All experiments were approved by the University of Memphis Institutional Animal Care and Use Committee.

### Drugs

ACEA, AA-5-HT, rimonabant, and dimethyl sulfoxide (DMSO) were obtained from Sigma-Aldrich (St. Louis, MO). All drugs were suspended in a vehicle containing 0.9% saline and 10% DMSO. ACEA (selective CB_1_R agonist) and AA-5-HT (dual FAAH/TRPV_1_ inhibitor) were each administered at 0.1, 0.5, and 1 mg/kg. Dose response curves for ACEA and AA-5-HT were constructed using 1 mg/kg as the highest dose, given that systemic 1 mg/kg doses of these drugs have been shown to modulate dopamine release in reward-related mesolimbic circuitry in rodents (Freels et al., 2019). The CB_1_R inverse agonist rimonabant was delivered at 0.3, 1, and 3 mg/kg, as rimonabant doses in this range have been shown to affect impulsive behavior in rats (Pattij et al., 2007).

### Apparatus

The RDT was conducted using MedAssociates (Fairfax, VA) operant chambers with retractable levers located to the left and right of an illuminable food trough. Chambers also included a house light, shock grating, pellet dispenser, and a 2.54 cm nose-poke port with recessed photobeams (0.635 cm). Behavior was reinforced using 45 mg sugar pellets (62.6% glucose, 26.8% fructose) obtained from Bio-Serv (Flemingon, NJ).

### Risky Decision-Making Task

Shaping and RDT procedures were conducted as previously described (Gabriel et al., 2019; Orsini et al., 2019). Initially, rats learned to associate the food trough with pellet delivery over a 60-minute session that included 38 individual pellet deliveries (inter trial interval [ITI] 100 ± 40 seconds) followed by food trough illumination. Food trough lights were extinguished after rats entered the trough to collect delivered pellets. Rats were then trained to press a single lever (left or right, counterbalanced across rats) on a fixed ratio-1 (FR-1) schedule to receive a single pellet reward. After rats achieved 50 presses on a single lever within 30 minutes, they then learned to press the opposite lever under the same criterion during the next training session. Upon completion of lever shaping, rats were trained to trigger extension of both levers by entering the food trough while it was illuminated. A single lever was extended with each trough entry in a pseudorandom order, with the same lever never being presented more than twice in a row. A lever press resulted in the delivery of 1 sugar pellet, and rats were trained until 35 presses on each lever was achieved during a session (70 total presses). One lever was then changed to a large-reward lever (3 pellets) while the other remained a small-reward lever (1 pellet). Trough entries then extended each lever 4 times in a pseudorandom order, for a total of 8 forced choice trials, followed by 10 choice trials wherein trough entries triggered extension of both levers. This 8 forced – 10 choice paradigm was repeated 5 times in a session for a total of 90 trials. Rats trained in this manner until they demonstrated discrimination between large- and small-reward levers (preference for the large-reward on more than 75% of trials, i.e. ~ 68 trials minimum). After rats completed all shaping, they began training in the RDT.

During RDT, rats were trained to choose between a small and large reinforcer accompanied by a variable risk of mild foot shock. Each session included 5 blocks of 18 trials (90 trials/session). Trial blocks began with 4 forced choice trials for each lever to establish punishment probabilities for large reward choices followed by 10 dual lever free choice trials. During forced choice trials, lever extension was pseudorandom such that the same lever was never presented more than 2 consecutive times. All trials started with illumination of both house and food trough lights, and rats had 10 seconds to enter the food trough. Trough entry extinguished the food trough light and resulted in the extension of one (forced choice trial) or both levers (free choice trial). Selection of the safe lever resulted in delivery of 1 pellet with 0% probability of foot shock, whereas selection of the risky lever triggered delivery of 3 pellets with a chance of a one second foot shock that increased across the 5 test blocks (0, 25, 50, 75, and 100% risk). After rewards were collected or 10 seconds passed (trial omission), house and food trough lights were extinguished and subsequent trials were initiated (ITI 10 ± 4 seconds, Figure 1A).

**Figure 1.**
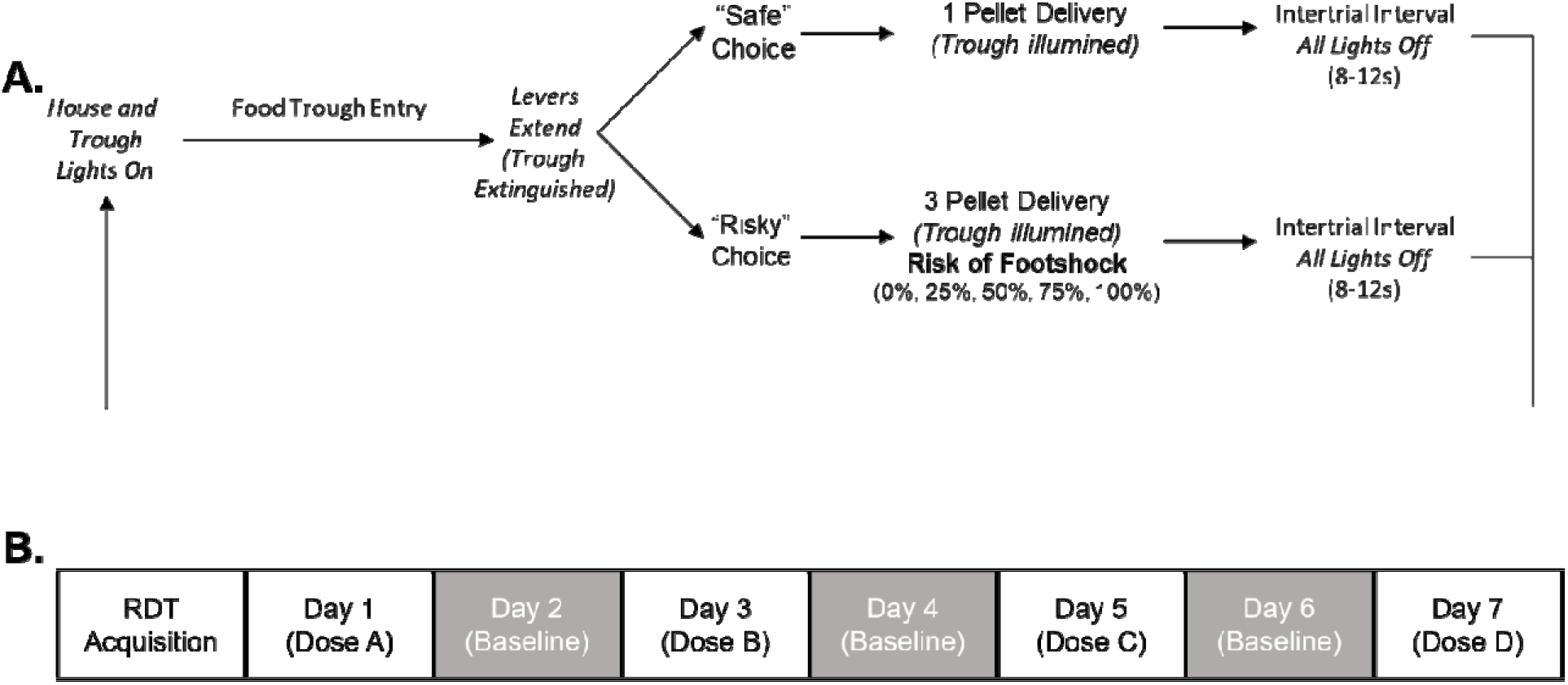
Risky decision-making task schematic and experimental timeline. (A) Visual representation of the Risky Decision-Making Task protocol adapted with permission from Gabriel et al. (2019). (B) Experimental timeline for pharmacological manipulations following acquisition of the Risky Decision-Making Task. Baseline days represent washout periods where no drugs were administered prior to behavioral testing.

RDT typically produces widespread variability, with some subjects demonstrating complete preference or avoidance of the risky reward (Simon et al., 2011). To induce a discounting curve while avoiding floor and ceiling effects, shock intensity was titrated within each rat. Foot shock amplitude began at 0.2 mA and was increased by either .02, .03, or .05 mA (due to variation in shock sensitivity) in the following session if rats completed greater than 85% of trials. After optimizing foot shock amplitudes, rats continued to train in the RDT until a stable discounting curve was maintained for a minimum of 3 days. Stability was assessed using a 2-way repeated measures ANOVA and was indicated by no main effect of test session and no test session x risk block interaction on the percentage of risky choices (Simon and Setlow, 2012).

### Drug Administration

Rats were systemically challenged with ACEA, AA-5-HT, rimonabant, or vehicle 30 minutes before RDT testing via intraperitoneal injection. A within-subjects design was used for the sequence of drug administration such that rats received their respective treatments on days 1, 3, 5, and 7 with baseline tests (no treatment) on days 2, 4, and 6 (Simon et al., 2011). Drug doses including vehicle administration were counterbalanced across treatment days (Figure 1B).

### Statistical Analysis

The impact of drug treatments on risk preference (% risky lever choices), choice latency (time to select a lever on free choice trials measured in seconds), and trial omissions were assessed. Analysis of risk preference was accomplished using 4 × 5 repeated measures ANOVAs with drug dose (vehicle, low, middle, high) and risk level (0, 25, 50, 75, 100% risk) as within subjects factors. Choice latency was investigated using 2 × 4 repeated measures ANOVAs with choice type (risky, safe) and drug dose as within subjects factors. Mean risk preference (i.e., % risky lever choices averaged across the 25, 50, 75, and 100% risk blocks) and omissions were analyzed using one-way repeated measures ANOVAs with drug dose as a within subjects factor. Significant main effects were further investigated using Bonferroni post-hoc tests, and interactions were probed using follow-up one-way repeated measures ANOVAs with Bonferroni post-hoc tests. All analyses were conducted using SPSS 23 (IBM Corp.).

## RESULTS

### Effects of Direct CB_1_R Activation on Risky Decision-Making

We first measured the effects of the direct CB_1_R agonist ACEA on risky decision-making. Analyses indicated a main effect of risk level across all doses (*F*_(4,32)_ = 25.773, *p* < 0.001, 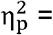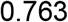), illustrating discounting of the large reward as probability of shock increased (Figure 2A). There was no main effect of ACEA dose (*F*_(3,24)_ = 0.252, *p* = 0.859, 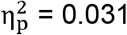) or dose x risk level interaction (*F*_(12,96)_ = 0.629, *p* = 0.813, 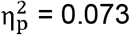) for ACEA on % risky lever choices (Figure 2A). Additionally, there was no main effect of ACEA dose on mean % risky lever choice across the entire session (*F*_(3,24)_ = 0.179, *p* = 0.910, 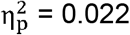; Figure 2B). Thus, acute CB_1_R activation does not alter punishment-based risky decision-making.

**Figure 2.**
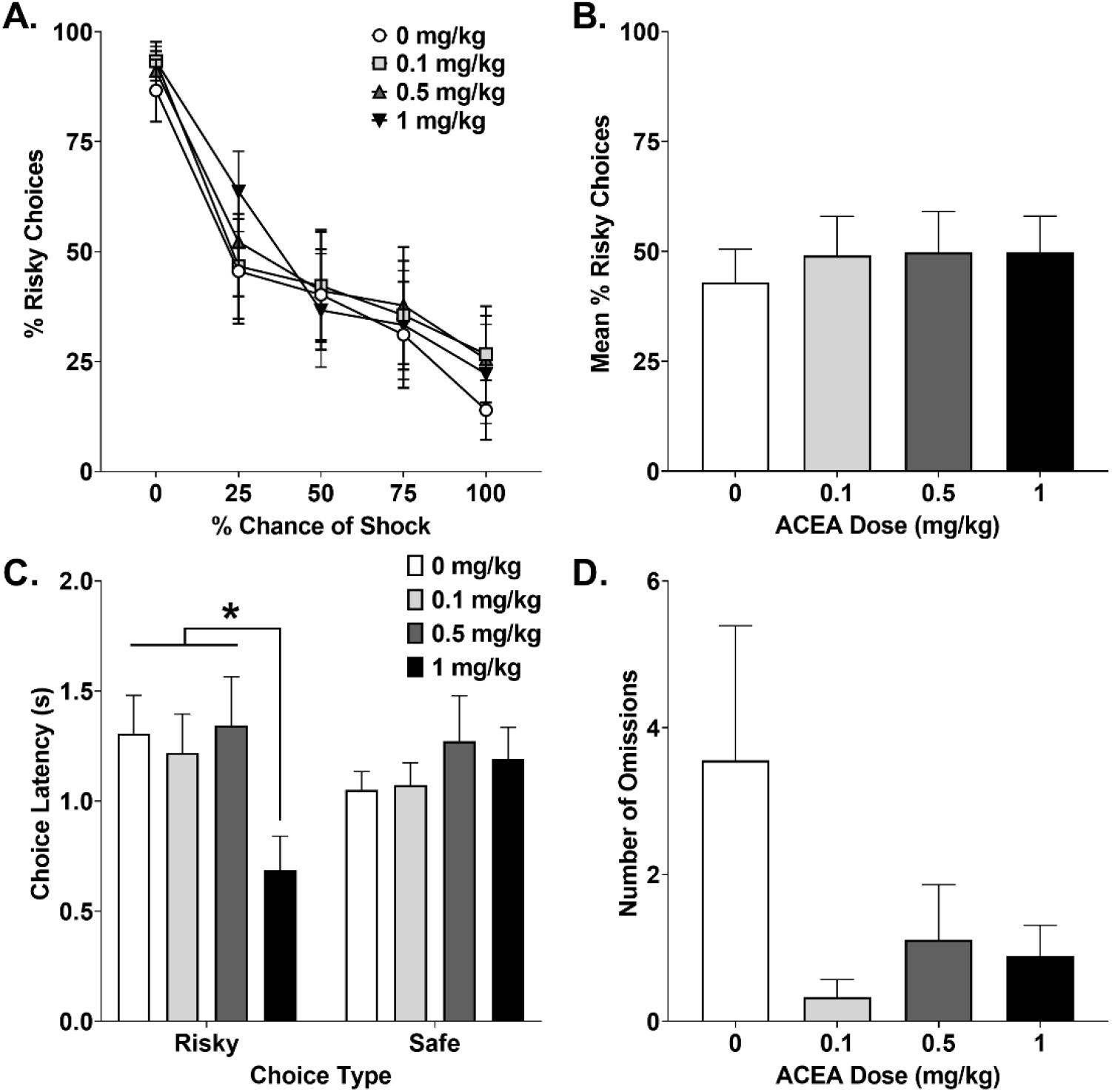
Effects of ACEA administration on performance in the Risky Decision-Making Task. (A) Risky reward discounting curves across blocks of increasing foot-shock probabilities. (B). Average % risky choices across test blocks that included a chance of foot shock. (C). Mean latencies for risky and safe decisions. (D) Total omissions committed during the Risky Decision-Making Task. * = *p* < 0.05. Error bars represent ± SEM.

Next, we measured the effects of direct CB_1_R activation on latency to make either a risky or safe decision. There was no main effect of choice type on choice latency (*F*_(1,8)_ = 0.002, *p* = 0.964, 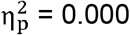), suggesting that rats used a comparable amount of time for deliberation to make risky or safe decisions. However, a significant ACEA dose x choice type interaction was revealed for choice latency (*F*_(3,24)_ = 8.977, *p* < 0.001, 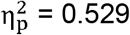), such that high dose ACEA reduced latency to make risky choices, but had no effect on safe choices (Figure 2C). Post-hoc analyses confirmed this observation, showing that risky choice latency was shorter at 1 mg/kg ACEA relative to 0, 0.1, and 0.5 mg/kg (*p*’s = 0.009 – 0.014), with no differences in safe choice latency between any doses (*p*’s = 0.502 – 1.000). There was no main effect of ACEA dose on trial omissions (*F*_(3,24)_ = 1.830, *p* = 0.169, 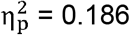; Figure 2D). Overall, CB_1_R activation reduced latency to make risky decisions, despite not affecting overall risk preference or task-engagement.

### Impact of CB_1_R Inverse Agonist Rimonabant on Risky Decision-Making

Next, we examined the effects of CB_1_R blockade with rimonabant on risky decision-making. As with direct CB_1_R agonism, there was a main effect of block on % risky lever choices (*F*_(4,36)_ = 13.054, *p* < 0.001, 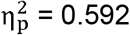), demonstrating that rats discounted the value of risky rewards across all drug doses. There was no main effect of rimonabant dose on % risky lever choice (*F*_(3,27)_ = 0.111, *p* = 0.953, 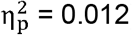), dose x risk level interaction (*F*_(12,108)_ = 1.579, *p* = 0.108, 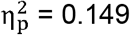), or effect of dose on mean % risky lever choices across the session (*F*_(3,27)_ = 0.110, *p* = 0.953, 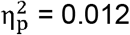; Figure 3A – B).

**Figure 3.**
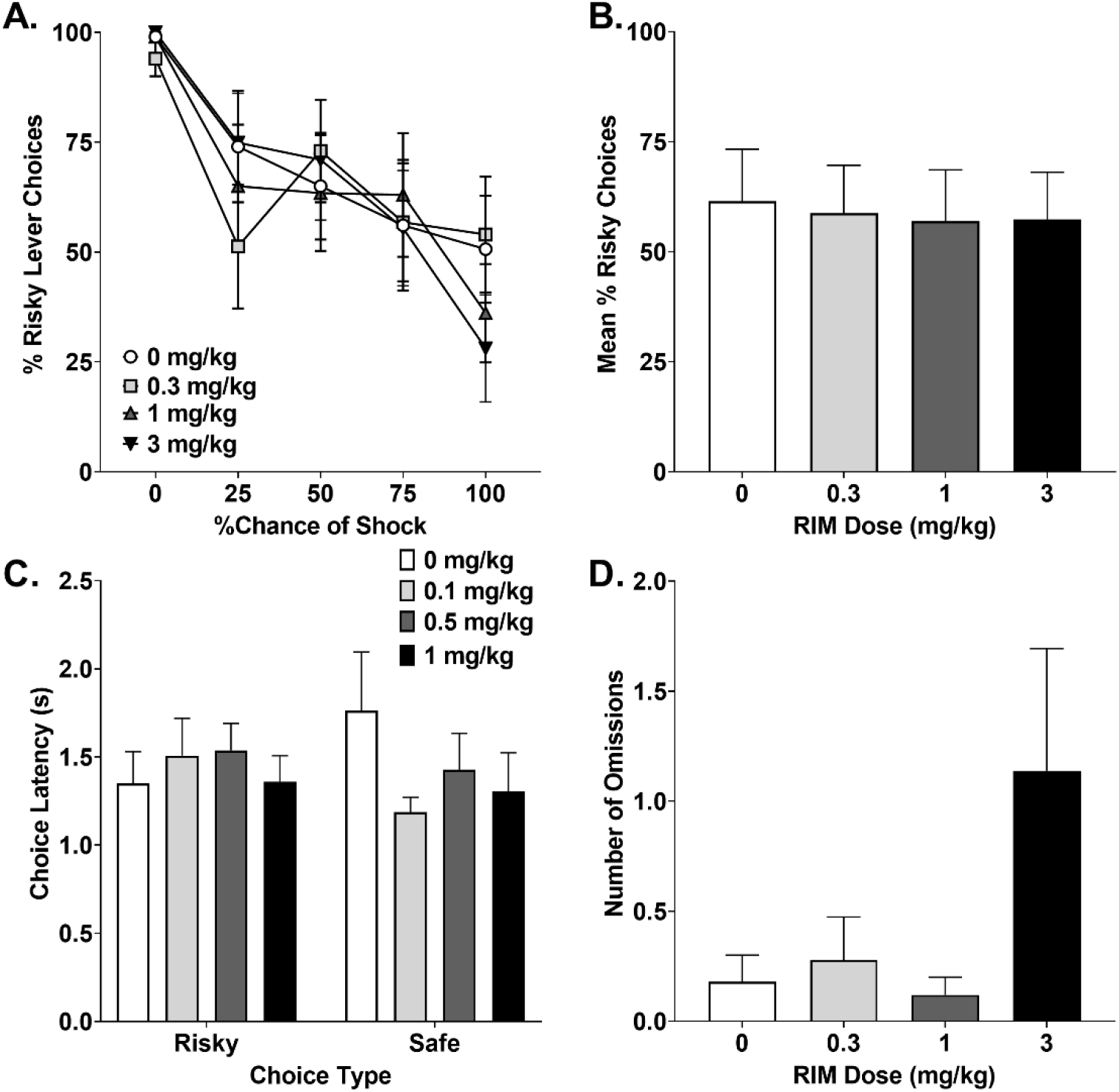
Influence of Rimonabant on Risky Decision-Making Task measures. (A) Mean % risky lever choices during test blocks. (B) Average % risky choices across test blocks with risk of foot-shock. (C) Risky and safe choice latencies. (D) Total omissions committed during testing. Error bars represent ± SEM.

We also tested the effects of CB_1_R blockade on decision-making latency. There was no difference in latency to choose risky vs safe options (*F*_(1,9)_ = 0.008, *p* = 0.932, 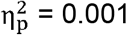). There was also no main effect of rimonabant dose on choice latency (*F*_(3,27)_ = 0.787, *p* = 0.512, 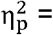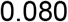), or dose x choice type interaction (*F*_(3,27)_ = 2.015, *p* = 0.135, 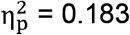; Figure 3 C). Additionally, analyses showed a trend towards a main effect of rimonabant dose on omissions (*F*_(3,27)_ = 2.801, *p* = 0.059, 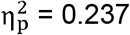, Figure 3D), with omissions only increasing with exposure to the high dose (3 mg/kg).

### Effects of the Indirect CB_1_R Agonist AA-5-HT on Risky Decision-Making

We measured the effects of indirect CB_1_R activation with AA-5-HT, a FAAH/TRPV1 inhibitor that increases overall endocannabinoid tone, on risky decision-making. As previously, a main effect of block on % risky lever choices was found (*F*_(4,32)_ = 9.823, *p* < 0.001, 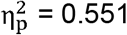) such that attenuation of risky choices occurred as probability of shock increased across all doses of drug. There was no effect of AA-5-HT dose on risk preference (*F*_(3,24)_ = 2.212, *p* = 0.113, 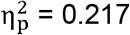; Figure 4A), dose x block interaction (*F*_(12,96)_ = 0.496, *p* = 0.912,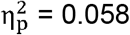), or effect of dose on mean % risky lever choices (*F*_(3,24)_ = 1.732, *p* = 0.187, 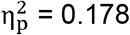; Figure 4B). Thus, similar to direct CB_1_R manipulation, AA-5-HT administration does not alter risk preference.

**Figure 4.**
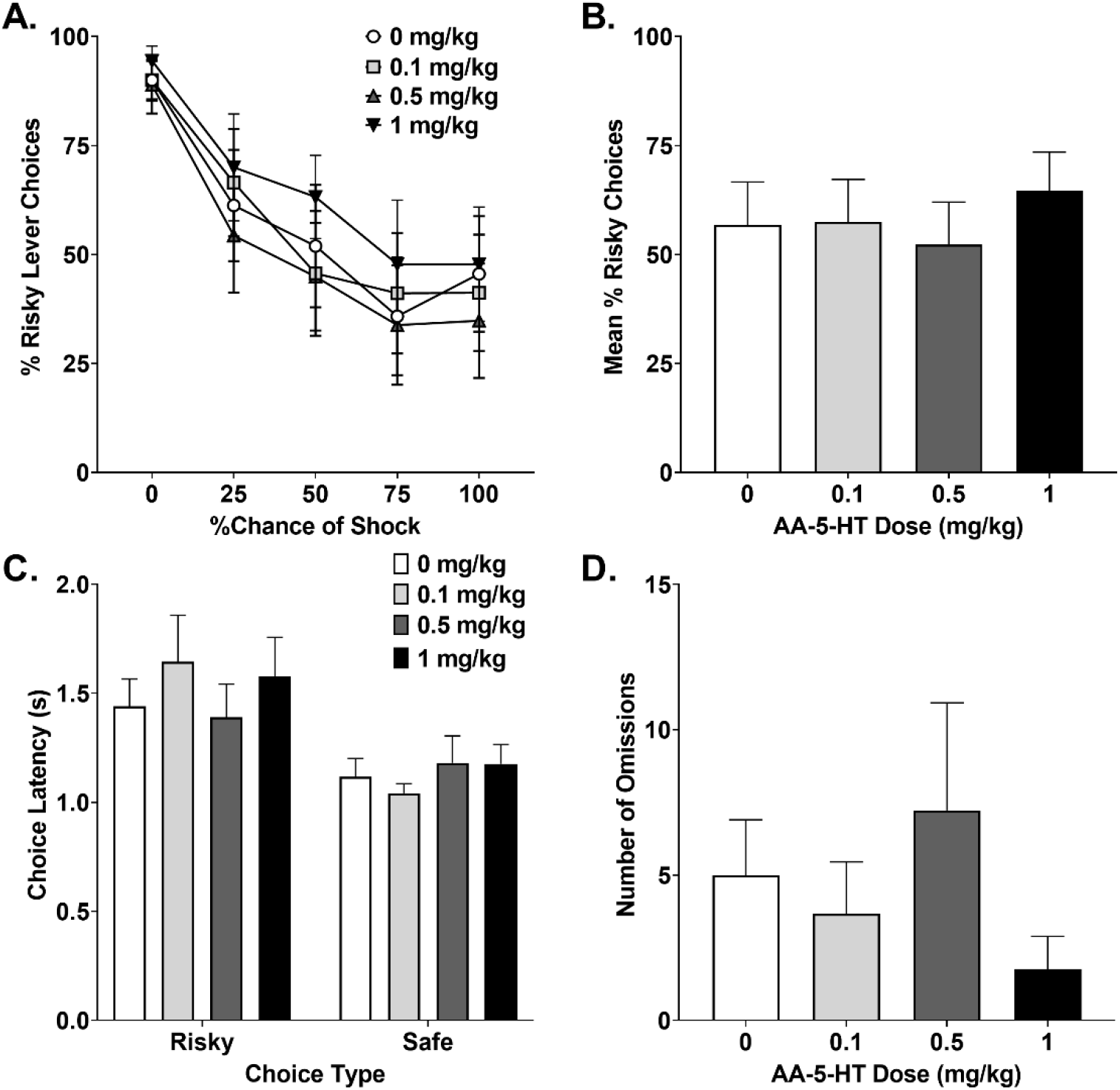
Risky Decision-Making Task performance following AA-5-HT treatment. (A) Average % risky choices within test blocks. (B) Mean % risky choices during test blocks with risk of foot-shock. (C) Choice latencies for risky and safe lever presses. (D) Total omissions committed during testing. Error bars represent ± SEM.

We next tested the effects of AA-5-HT on decision-making latency. There was a main effect of choice type on choice latency (*F*_(1,8)_ = 8.809, *p* = 0.018, 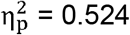) such that rats were slower to make risky choices than safe choices (Figure 4C). Individual comparisons between safe and risky choice latency at each drug dose revealed that this difference was only significant at the low dose of .1mg/kg (*p* = .015), but not with saline or higher doses (*p* >.05). There was no overall main effect of dose on latency (*F*_(3,24)_ = 0.355, *p* = 0.786, 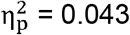) or dose x choice type interaction (*F*_(3,24)_ = 1.538, *p* = 0.230, 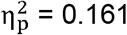; Figure 3C). There was also no main effect of AA-5-HT dose on omissions(*F*_(3,24)_ = 2.077, *p* = 0.130, 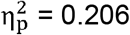: Figure 4D).

## DISCUSSION

Given the rise in permissive cannabis use policies and the lack of understanding regarding the effects of cannabinoids on cognitive function, we investigated the contribution of CB_1_R activity to risk-based decision making. We measured risk-taking with the RDT, in which rats choose between small and large rewards associated with risk of foot shock. Our results indicated that bidirectional CB_1_R manipulation does not affect risky decision-making. However, direct CB_1_R activation, but not CB_1_R blockade or indirect CB_1_R agonism, selectively reduced latency to make risky decisions while not affecting safe choice latency. Together, our results suggest that CB_1_Rs are not directly involved with valuation of rewards and risk of punishment during decision-making, but are able to accelerate risky decisions.

### CB_1_R Activation does not Affect Punishment-Based Risk-Taking

In the RDT, we found that direct CB_1_R agonism, indirect CB_1_R stimulation, and CB_1_R blockade did not influence overt risk-taking behavior, as indicated by the lack of effect of drug treatment on the preference for rewards associated with risk of punishment. Although there is little preclinical research regarding the effects of cannabinoids on risky decision-making, our results corroborate findings obtained using the rat gambling task (rGT), in which rats choose between rewards of different magnitude and probability with failed trials resulting in a time-out punishment (Zeeb et al., 2009). Administration of THC, synthetic CB_1_R agonists, indirect CB_1_R agonists, or CB_1_R antagonists did not impact risk-taking in the rGT, apart from direct CB_1_R agonism improving choice strategy in suboptimal rats only (Gueye et al., 2016; Ferland et al., 2018). However, human research has revealed that acute and chronic THC exposure elicits increased risk-taking in gambling tasks depending on the individual’s history of cannabis use (Lane et al., 2005; Vadhan et al., 2007; Fridberg et al., 2010). It is important to note that the RDT is distinct from preclinical and clinical gambling models in that it utilizes physical punishment as a negative outcome of risk-taking rather than reward omission or loss. This modality of punishment captures a distinct form of risk taking reliant upon different neural mechanisms than omission-based risk tasks (see Orsini et al., 2015a for review). Contrasting results obtained from the RDT and gambling-like procedures may indicate that the neural mechanisms recruited during these tasks exhibit differential sensitivity to CB_1_R manipulation. Therefore, our results extend current research by suggesting that while CB_1_Rs may be involved in modulating some forms of risky decision-making, this may not translate to decision-making under risk of physical consequences.

### Direct CB_1_R Activation Reduces Risky Choice Latency

Although we found no evidence for CB_1_R modulation of risk-preference in the RDT, we observed that high dose direct CB_1_R agonism reduced the latency for rats to make risky decisions. Interestingly, CB_1_R mediation of decision-making rate did not occur during safe choices with no risk of consequences. Thus, systemic CB_1_R activation reduces the time an organism spends weighing the perceived value of a reward against the cost of possibly experiencing an adverse outcome following risk-taking. One potential explanation for this increased behavioral efficiency is that CB_1_R agonism attenuates the aversive emotional state associated with impending punishment. Systemic CB_1_R activation in humans has been associated with attenuated anxiety and stress, and produces anti-fear effects (Rabinak et al., 2013; Childs et al., 2017; Cuttler et al., 2018), and complementary data from rodent models demonstrate that CB_1_R agonist treatments have been associated with reduced aversive learning, impaired fear memory acquisition, enhanced fear extinction learning, reduced stress responsivity, and anxiolysis (Kangarlu-Haghighi et al., 2015; Simone et al., 2015; Nasehi et al., 2016; Fokos and Panagis, 2010; Gobira et al., 2013; Kangarlu-Haghighi et al., 2015; Kinden and Zhang, 2015; Schreiber et al., 2018; Uttl et al., 2018). Interestingly, the suppressive effects of cannabinoids on anxiety, fear, and stress appear to stem from CB_1_R activation in the basolateral amygdala (BLA), which is also involved in the integration of information regarding risk of punishment and reward value (Ganon-Elazar and Akirav, 2009; Orsini et al., 2015b; Morena et al., 2016). Therefore, systemic CB_1_R activation may have altered BLA functioning during anticipation of risky choice, thereby attenuating anxiety-, fear-, and/or stress-related processes associated with risk of punishment that typically interrupt motivated behavior (Kim et al., 2014).

Alternatively, it may be that the accelerated risky choices are related to impulsive action, defined as the inability to inhibit a previously beneficial response (Bari and Robbins, 2013). CB_1_R activation has been shown to elevate impulsive action (Ramaekers et al., 2006; Irimia et al., 2015; Ferland et al., 2018), which may drive rats to select risky options quickly without deliberation about potential punishment, especially considering the lack of punishment in the first block of the task. However, lack deficit in inhibitory control would also be expected to reduce the ability to shift away from the risky reward with increasing probability of punishment, which was not observed here. Another possibility for the elevated rate of risk-taking is that CB_1_R activation caused antinociception that diminished the ability of the impending shock to affect behavior (Bridges et al., 2001; Wibelhaus et al., 2012). However, it is unlikely that the effects of direct CB_1_R stimulation on risky choice latency were solely due to gross effects on pain tolerance given indirect CB_1_R activation (which also has analgesic effects) did not affect choice latencies (Maione et al., 2007; de Novellis et al., 2008). Furthermore, research has indicated that sensitivity to punishment is not related to risk preference in the RDT (Simon et al., 2011). Another potential explanation is that CB_1_R activation may influence motivation to obtain risky rewards, as previous studies have indicated that shorter response latencies are associated with increased incentive value of rewards (Holland and Straub, 1979). Additionally, CB_1_Rs have been implicated in the facilitation of motivated responding for rewards (Ward and Dykstra, 2005; Maccioni et al., 2008; Helfand et al., 2017). Thus, CB_1_R activation may selectively enhance the motivational properties of risky rewards while not influencing overall motivation in the RDT, indicated by the absence of effects on safe reward choice latency or trial omissions.

In contrast to direct CB_1_R agonism, indirect CB_1_R agonist treatments did not influence risky choice latency relative to vehicle treated rats. This discrepancy could be due in part to differences in pharmacological actions between ACEA and AA-5-HT. While ACEA is a highly selective CB1R agonist, AA-5-HT is a dual FAAH/TRPV1 blocker (Hillard et al., 1999; Maione et al., 2007). Therefore, non-specific effects of AA-5-HT could have negated the reduction in risky choice deliberation. Another option is that acute systemic administration of AA-5-HT within the dose ranges used may not have affected the negative emotional responses potentially associated with punishment following risky decisions. Previous research has indicated that the anxiolytic effects of AA-5-HT are dose-, rodent strain-, and context- dependent, and in some cases chronic administration regimens are required to elicit anxiolysis (Micale et al., 2009a,b; John and Currie, 2012; Freels et al., 2019). Additionally, the influence of AA-5-HT on fear has been found to be dependent on the type of fear conditioning (i.e., auditory vs. contextual fear conditioning) and rodent strain used (Gobira et al., 2017; Freels et al., 2019). Furthermore, the attenuation of the neuroendocrine response to stress by AA-5-HT is also dose-dependent and appears to manifest following multiple AA-5-HT treatments (Navarria et al., 2014). Therefore, future experiments using a wider range of AA-5-HT doses as well as chronic treatment regiments are required to fully assess whether AA-5-HT influences aspects of risky decision-making within the context of the RDT.

Similar to indirect CB_1_R agonism, no changes in risky choice latencies were observed following CB_1_R blockade, suggesting that CB_1_Rs do not control risk-taking latency in a bidirectional manner. In contrast, results obtained from the rGT have shown that CB_1_R antagonist administration within the dose range that was used here in the RDT increases choice latency (Ferland et al., 2018). These differences may stem from the differences in task parameters between the rGT and RDT such as punishment modality. However, it should be noted that CB_1_R antagonism with AM 4113 has also been found to not influence choice latency in the rGT (Gueye et al., 2016). Given that CB_1_R antagonism did not influence choice latency (or risk preference) in the RDT, it is possible that CB_1_Rs are not necessary for driving punishment-based risky decision-making, but may specifically influence the deliberation of risk and reward outcomes.

### Summary and Conclusions

In summary, CB_1_R manipulation dose not influence risk preference in punishment-based risky decision-making, but selectively increases the speed of risky decisions. Although overall risky decision-making is not affected by CB_1_R activation, the lack of hesitation to make risky choices could lead to “rushed” decisions to engage in risk-taking despite potential adverse consequences in human cannabis users. Additionally, these experiments suggest that risky decision-making and rate of deliberation have distinct pharmacological substrates, which is an important consideration for the development of precise treatments for risky decision-making deficits commonly observed in SUDs. In conclusion, the potential contribution of systemic CB_1_R activation to the development of SUD-related impairments in risk-taking may stem from subtle alterations in punishment-based risky decision-making and more prominent impairments in forms of risk-taking that are related to probabilistic discounting of risky rewards.

